# Genome of *Solanum pimpinellifolium* provides insights into structural variants during tomato breeding

**DOI:** 10.1101/2020.06.17.157859

**Authors:** Xin Wang, Lei Gao, Chen Jiao, Stefanos Stravoravdis, Prashant S. Hosmani, Surya Saha, Jing Zhang, Samantha Mainiero, Susan R. Strickler, Carmen Catala, Gregory B. Martin, Lukas A. Mueller, Julia Vrebalov, James J. Giovannoni, Shan Wu, Zhangjun Fei

**Author notes:** These authors contributed equally (Xin Wang, Lei Gao). Correspondence should be addressed to Zhangjun Fei and Shan Wu.

## Abstract

*Solanum pimpinellifolium* (SP) is the wild progenitor of cultivated tomato. Because of its remarkable stress tolerance and intense flavor, SP has been used as an important germplasm donor in modern breeding of tomato. Here we present a high-quality chromosome-scale genome sequence of SP LA2093. Genome comparison identifies more than 92,000 high-confidence structural variants (SVs) between LA2093 and the modern cultivar, Heinz 1706. Genotyping these SVs in ~600 representative tomato accessions unravels alleles under selection during tomato domestication, improvement and modern breeding, and discovers numerous novel SVs underlying genes known to regulate important breeding traits such as fruit weight and lycopene content. Expression quantitative trait locus (eQTL) analysis detects hotspots harboring master regulators controlling important fruit quality traits, including cuticular wax accumulation and flavonoid biosynthesis, and novel SVs contributing to these complex regulatory networks. The LA2093 genome sequence and the identified SVs provide rich resources for future research and biodiversity-based breeding.

## Introduction

Tomato (*Solanum lycopersicum*) is the world’s leading vegetable crop with a total production of 182 million tons and a worth over US$60 billion in 2018 (http://www.fao.org/faostat). *S. pimpinellifolium* (SP) carrying red, small and round fruits is the wild progenitor of the cultivated tomato. It was domesticated in South America to give rise to *S. lycopersicum* var. *cerasiforme* (SLC), which was later improved into the big-fruited tomato *S. lycopersicum* var. *lycopersicum* (SLL) in Mesoamerica (Blanca et al., 2012; Blanca et al., 2015). The fact that SP can freely hybridize with SLL has enabled the incorporation of SP alleles into modern tomato cultivars to improve disease resistance, abiotic stress tolerance and other fruit quality traits (Ebert and Schafleitner, 2015). Due to the importance of SP, draft genome assemblies of two accessions, LA1589 (Tomato Genome Consortium, 2012) and LA0480 (Razali et al., 2018), have been generated using Illumina short-read sequencing technology. Although these assemblies provided some essential genomic information for SP, they are highly fragmented and incomplete, limiting their applications as robust references in tomato breeding and research.

Genomic structural variants (SVs), including insertions/deletions (indels), inversions and duplications, are the causative genetic variants for many domestication traits of crops (Gaut et al., 2018). Empirical cases in tomato include a ~294-kb inversion at the *fas* locus leading to enlarged fruits (Xu et al., 2015), a 1.4-kb deletion in the *CSR* gene resulting in an increased fruit weight (Mu et al., 2017), a partial deletion of the *LNK2* gene leading to the circadian period lengthening in cultivated tomatoes (Muller et al., 2018), and a tandem duplication at *sb3* that suppresses the excessive inflorescence branching (Soyk et al., 2019). Whole-genome SNP data have been employed to reveal the impact of human selection on the tomato genome (Lin et al., 2014) and reconstruct tomato domestication history (Razifard et al., 2020). However, the distribution of SVs in tomato accessions and their population dynamics are largely unexplored. Identification of SVs between SP and SLL, studying their evolutionary dynamics in different tomato populations and investigating their regulatory roles in the context of gene expression can provide critical insights into the contribution of SVs to important agronomic traits in tomato domestication and breeding.

In this study, we present a high-quality chromosome-scale genome sequence of an SP accession, LA2093, assembled from PacBio long reads combined with Hi-C chromatin interaction maps. LA2093 harbors many desirable traits and has served as the donor parent of a recombinant inbred line (RIL) population that has been widely used for mapping disease resistance and fruit quality traits (Ashrafi et al., 2009; Ashrafi et al., 2012; Kinkade and Foolad, 2013; Gao et al., 2019; Gonda et al., 2019). We further identified high-confidence SVs between the genomes of LA2093 and cultivar Heinz 1706 through direct genome comparison combined with PacBio long read mapping, and genotyped this reference set of SVs in ~600 tomato accessions representing SP, SLC, and heirloom and modern SLL to determine their population dynamics. We also employed expression quantitative trait locus (eQTL) mapping to explore the roles of SVs in regulating gene expression and identified distant-acting eQTL hotspots and potential master regulators controlling important fruit traits.

## Results

### Sequencing and assembly of the *S. pimpinellifolium* genome

We assembled the genome of LA2093 using PacBio long reads and Hi-C chromatin contact information. A total of 96 Gb of PacBio sequences with an N50 read length of 22.3 kb was generated, covering approximately 103× of the LA2093 genome with an estimated size of 923 Mb (**Supplementary Fig. 1**). The PacBio reads were *de novo* assembled into contigs, followed by polishing with both PacBio and Illumina reads. This resulted in an assembly of 453 contigs with a total length of 807.6 Mb and an N50 length of 10.9 Mb (**Supplementary Table 1**). A total of 166 million Hi-C read pairs was generated for constructing chromatin interaction maps. These Hi-C contact maps, together with the synteny with the Heinz 1706 genome (version 4.0) (Hosmani et al., 2019) and the genetic map constructed using the NC EBR-1 × LA2093 RIL population (Gonda et al., 2019), were used to scaffold the assembled contigs. Finally, 385 contigs with a total length of ~800 Mb, accounting for 99.0% of the assembly, were clustered into 12 pseudomolecules (**Fig. 1a**). The Hi-C heatmap (**Supplementary Fig. 2**) and the good collinearity between the pseudomolecules and the genetic map (**Supplementary Fig. 3**) supported the chromosome-scale structure of the assembly. BUSCO (Simao et al., 2015) assessment indicated that about 97.8% of the core conserved plant genes were found complete in the LA2093 assembly (**Supplementary Table 1**). Collectively, these results indicated that the LA2093 genome assembly is of high quality, with substantially improved contiguity and completeness compared to the two previously reported *S. pimpinellifolium* genome assemblies (Tomato Genome Consortium, 2012; Razali et al., 2018) (**Supplementary Table 1**). A total of 544.3 Mb repetitive sequences (67.3%) were identified in the LA2093 assembly with the *gypsy* retrotransposon being the most abundant repeat family (37.8%; **Supplementary Table 2**). The genome was predicted to harbor 35,761 protein-coding genes, of which 35,535 (99.4%) were supported by RNA-Seq data, and/or homologs in the NCBI non-redundant protein database.

**Fig. 1.**
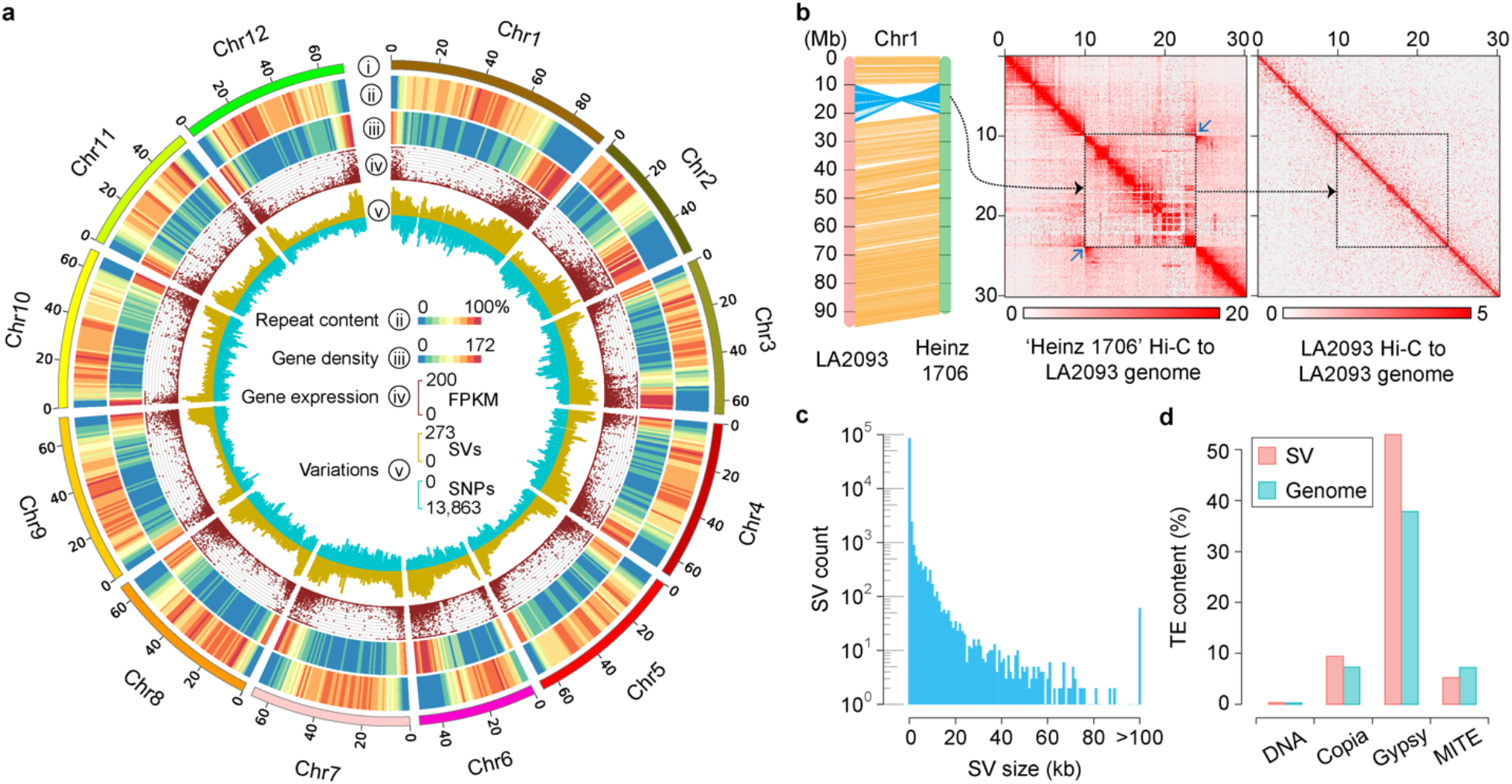
Genomic landscape of *S. pimpinellifolium* LA2093 and structural variants (SVs) identified between LA2093 and Heinz 1706. **a**, Features of the LA2093 genome. i, Ideogram of the 12 chromosomes in Mb scale. ii, Repeat content (% nucleotides per Mb); iii, Gene density (number of genes per Mb). iv, Gene expression (FPKM). v, Densities of SVs (outer) and SNPs (inner) in comparison to Heinz 1706 (number of SVs and SNPs per Mb). **b**, Alignment of chromosome 1 between LA2093 and Heinz 1706. The color intensity in Hi-C heatmaps represents the number of links between two 100-kb windows. The inversion shown in blue (left) is supported by high-density contacts pointed by the two blue arrows in Hi-C heatmaps generated from Heinz 1706 Hi-C reads aligned to the LA2093 genome (middle), while no corresponding contract is found in the LA3093 Hi-C heatmap (right). **c**, Distribution of SV sizes. **d**, Contents of different categories of transposable elements in SV regions and the whole genome of LA2093.

### Genomic SVs between LA2093 and Heinz 1706

Alignment of the genome sequences of LA2093 and Heinz 1706 (version 4.0) (Hosmani et al., 2019) showed good collinearity between the two reference genomes (**Supplementary Fig. 4**). Despite the high collinearity, 28 inversions ranging from 483 bp to 13.9 Mb were identified distributing across all 12 chromosomes **(Supplementary Table 3**), which were further supported by PacBio reads and/or Hi-C maps (**Fig. 1b** and **Supplementary Fig. 5)**. Together these inversions harbored approximately 800 genes (861 in LA2093 and 733 in Heinz 1706), of which a large portion (457) were annotated with functions related to disease resistance and response to abiotic stress (**Supplementary Table 4**). We detected significantly lower diversity of both indels and SNPs inside these inversions (**Supplementary Fig. 6**), consistent with the assumption that preexisting ancient inversions may cause genetic diversity departure from neutrality (Guerrero et al., 2012). These inversions could cause linkage drag and recombination restriction. Indeed, restriction of recombination was detected around the tomato fruit weight QTL, *fw1.1* (12.5-20.7 Mb on chromosome 1) (Illa-Berenguer et al., 2015), which could be associated with the large inversion identified on chromosome 1 from 9.3 to 20.4 Mb (**Fig. 1b**). Our results here provide information for more efficient breeding and QTL fine-mapping through inversion-aware approaches.

In addition to inversions, indels between LA2093 and Heinz 1706 genomes were detected by direct comparison of the two high-quality assemblies combined with mapping of PacBio long reads to the genomes. A total of 92,522 high-confidence indels, ranging from 10 bp to 2.4 Mb, were identified (**Fig. 1a, c** and **Supplementary Table 5**). As expected, the majority of the indels were relatively short with 82.8% <100 bp and only 0.1% >100 kb (**Fig. 1c**). Approximately 52.9% of these indel sequences were *gypsy*-like retrotransposons, compared to 37.8% of the entire genome, while the contents of other types of transposable elements were similar between the indel regions and the whole genome (**Fig. 1d**), suggesting that indels occurred more frequently in genome regions occupied by *gypsy*-like retrotransposons.

SVs in gene body and promoter regions can directly impact gene functions and expression. Only 14.8% of identified indels overlapped with gene body and promoter (defined here as 3-kb upstream of gene body) regions, notably lower than the proportion in the whole genome (27.7%), implying a functional constraint against these indels on coding or regulatory regions **(Supplementary Table 6)**. More than half of the predicted genes in LA2093 (21,875 out of 35,761) and Heinz 1706 (19,590 out of 34,689) were affected by at least one indel in their gene body or promoter regions. Genes affected by the identified indels were enriched with those involved in response to stimulus, reproduction, signal transduction, and primary and secondary metabolic processes (**Supplementary Fig. 7**), suggesting that these indels may contribute to the differences in disease resistance and fruit quality traits between the wild and cultivated tomatoes. We detected several SVs known to underlie tomato domestication traits, such as the 1.4-kb deletion and 22-bp insertion in the *CSR* gene leading to increased fruit weight (Mu et al., 2017), the 7-kb deletion in the *LNK2* gene responsible for the circadian period lengthening in cultivated tomatoes (Muller et al., 2018), the 4-kb substitution in the promoter of the *TomLoxC* gene, which contributes to fruit flavor (Chen et al., 2004; Shen et al., 2014; Gao et al., 2019), and the 85-bp deletion in the promoter of the *ENO* gene, which regulates floral meristem activity (Yuste-Lisbona et al., 2020) (**Supplementary Table 7**).

### Selection of SVs in tomato domestication and breeding

*De novo* detection of SVs based on short read alignments to a reference genome is subject to a relatively high rate of both false negatives and false positives (Handsaker et al., 2011; Mills et al., 2011). Therefore, the high-confidence SVs identified between LA2093 and Heinz 1706 through direct genome comparison and PacBio long read mapping provided a valuable set of reference SVs that could be used to investigate the roles of SVs in tomato domestication and breeding. The 92,522 indels and 28 inversions were genotyped in 597 tomato accessions, including 51 SP, 6 *S. cheesmaniae* and *S. galapagense* (SCG), 228 SLC, 226 heirloom, 52 modern and 34 other cultivars (**Supplementary Table 8**). To estimate the accuracy of SV genotyping using short read mapping to the reference SVs, we generated 31.1 Gb of Nanopore long read sequences for an SP accession, LA1589, and aligned the long reads to the LA2093 and Heinz 1706 genomes for SV calling. About 96.7% of the genotypes determined using the LA1589 short reads were confirmed by the Nanopore long read mapping method, indicating a high reliability of our SV genotyping. Phylogenetic and population structure analyses using the SVs clearly separated the SP and SCG groups from the heirloom and modern groups with SLC being the intermediate group between the wild and cultivated accessions. A similar pattern was observed using whole-genome SNPs (**Supplementary Fig. 8**), further assuring the reliability of our identified SVs. Seven accessions positioned into unexpected species groups were excluded from downstream analyses (**Supplementary Table 8**).

SV allele frequency changes among different tomato populations are a result of evolutionary events, such as selection of desirable traits, reduced population size, and introgression from ancestral groups. To identify SVs under selection during tomato domestication and breeding, we investigated SV allele frequency changes from SP to SLC for domestication, from SLC to SLL heirlooms for improvement, and from heirlooms to modern elite lines for modern breeding. In the SP population, the homozygous LA2093 alleles of the SVs were prevalent, taking up an average of 58.4% of the SVs in each accession, while only 39.4% SVs were homozygous for the Heinz 1706 genotypes (**Fig. 2a**). After domestication and improvement, the frequencies of Heinz 1706 alleles increased to 84.3% and 95.3% in SLC and heirloom tomatoes, respectively, and then slightly decreased to 93.6% in modern lines (**Fig. 2a**). These findings implied significant genetic diversity loss imposed by domestication, especially the loss of the SP-specific alleles. The allele frequencies of 38,367 SVs were significantly changed between different tomato populations (**Supplementary Table 5**). During domestication, the LA2093 allele frequencies of 17 inversions and 37,632 indels were significantly lower in SLC than in SP, while only 217 indels had the LA2093 allele frequencies significantly higher in SLC (**Fig. 2b**). These selected indels could affect 14,189 genes in LA2093 and 12,264 in Heinz 1706. In the improvement process, only 103 indels had higher LA2093 allele frequencies in heirlooms than in SLC, and 25,579 SVs (13 inversions and 25,566 indels) displayed significantly lower LA2093 allele frequencies in heirlooms (**Fig. 2b**), which collectively could affect 9,530 and 7,870 genes in LA2093 and Heinz 1706, respectively. The enriched functions were shared between genes affected by the SVs selected during domestication and improvement, including stress and stimulus response, biosynthesis, cell differentiation, embryo development, pollination and reproduction processes (**Fig. 2c,d**). These results demonstrated a common selection preference for the Heinz 1706 alleles in tomato domestication and improvement. It is worth noting that, despite the continuous loss of wild species alleles from SP to SLC and SLL, 1,397 SVs exhibited significantly higher LA2093 allele frequencies in modern lines than in heirlooms (**Fig. 2b**). This could be related to the re-introduction of agriculturally favorable alleles from wild accessions into modern SLL lines.

**Fig. 2.**
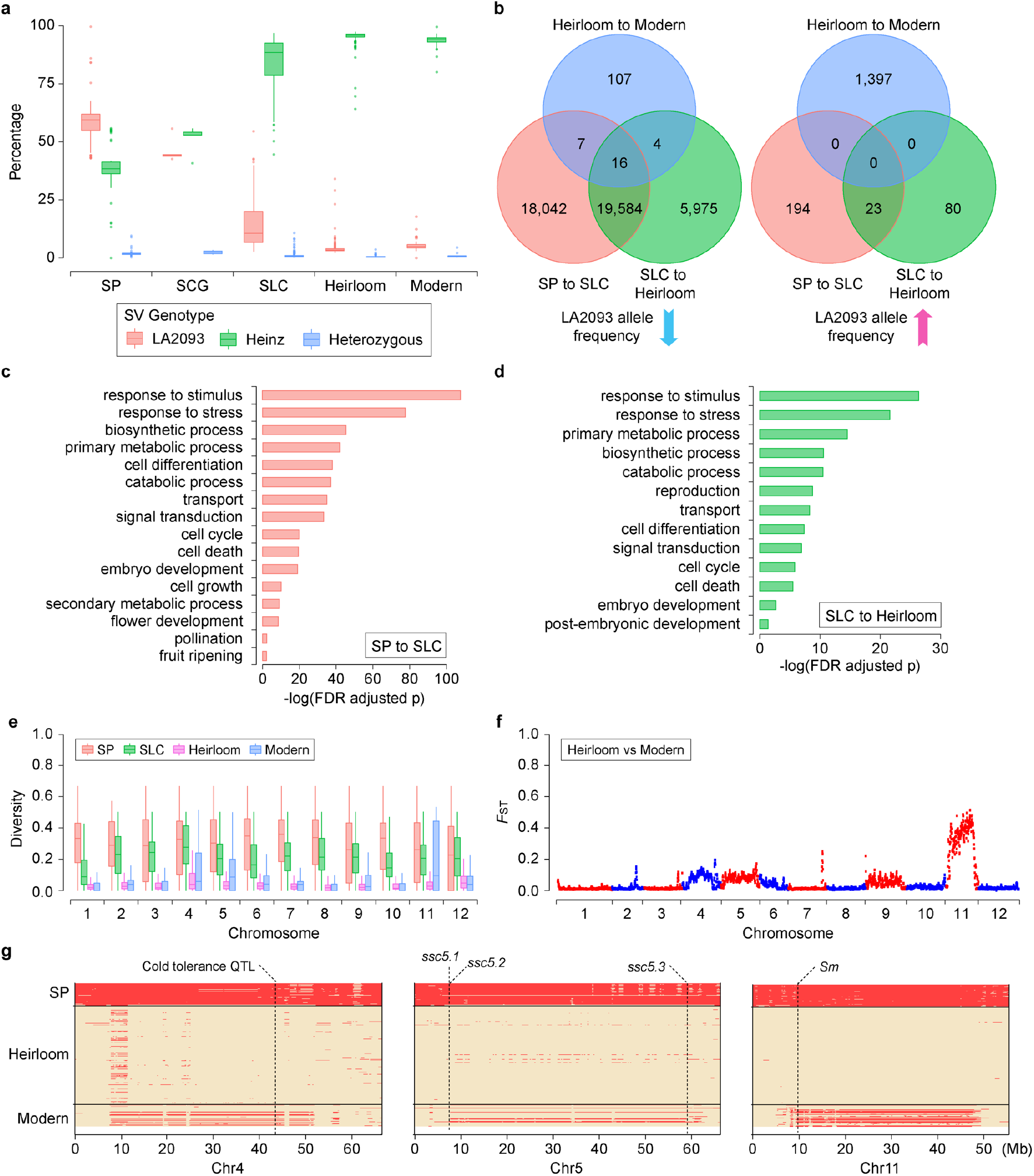
SVs under selection during tomato domestication and breeding. **a**, Percentages of SVs with different genotypes in each accession of different groups. **b**, Venn diagrams of selected SVs during domestication (SP to SLC), improvement (SLC to heirloom) and modern breeding (heirloom to modern). **c-d**, GO terms enriched in genes affected by SVs selected during domestication (**c**) and improvement (**d**). **e**, SV diversity in different groups across the 12 chromosomes. **f**, Distribution of *F*_ST_ values between heirloom and modern groups across the LA2093 genome. **g**, Introgressions from SP to modern tomatoes. Known QTLs and genes are indicated by the dash lines. For each box plot, the lower and upper bounds of the box indicate the first and third quartiles, respectively, and the center line indicates the median.

The diversity of SVs in the modern group was similar to that of the heirlooms but lower than that in SLC and SP for most genomic regions. However, a substantially higher SV diversity was observed in the modern group compared to the heirlooms on chromosomes 4, 5 and 11 (**Fig. 2e**), consistent with the higher *F*_ST_ values between the two populations on these chromosomes (**Fig. 2f**). Concordantly, introgression of genomic regions from SP to modern cultivars were identified in all these three chromosomes (**Supplementary Fig. 9**). The introgressions on chromosome 4 contained a cold tolerance QTL (Foolad et al., 1998), on chromosome 5 carried several known QTLs controlling soluble solid content (SSC) in fruit (Lin et al., 2014), and on chromosome 11 included an important disease resistant genes, *Sm* (Su et al., 2019) (**Fig. 2g**). These results imply that SP introgressions in the modern cultivars might be acquired through contemporary breeding to introduce abiotic and biotic stress tolerance and favorable flavor traits.

### SV selection associated with breeding traits

Tomato fruit phenotypes have changed dramatically during domestication and improvement. Investigating population dynamics of SVs potentially affecting the expression or functions of genes controlling important horticultural traits can improve our understanding of the impacts of human selection on these genes and provide potential targets for breeding. Fruit size enlargement is a major domestication syndrome in tomato. Several SVs have been identified to be associated with fruit weight, including the 1.4-kb deletion in *CSR* and the 85-bp deletion in the promoter of *ENO*, both of which underlie larger fruit size (Mu et al., 2017; Yuste-Lisbona et al., 2020). Consistent with that reported in Mu et al. (2017), we found that the allele frequency of the 1.4-kb deletion in *CSR* was 1.0% and 11.5% in SP and SLC, respectively, and became largely fixed in SLL with 90.2% in heirlooms and 94.1% in modern lines (**Supplementary Table 7**). The 85-bp deletion in the *ENO* promoter had a frequency of 54.7% in SP, 90.8% in SLC, 94.0% in heirloom and 96.8% in modern accessions. Interestingly, we identified an additional 257-bp insertion in *CSR* spanning the 5’ UTR and the CDS, which had a frequency of 38.6% in SP, 77.9% in SLC, 98.3% in heirloom and 97.7% in modern accessions, and another 57-bp deletion in the *ENO* promoter, whose allele frequencies in SP, SLC, SLL heirloom and modern were 22.0%, 83.5%, 92.3% and 94.6%, respectively. Furthermore, novel indels were detected for additional fruit weight genes, including a 23-bp insertion in the promoter of *RRA3a* and a 14-bp deletion in the promoter and a 12-bp insertion in the first exon of *CLE9* (Xu et al., 2015), all of which had allele frequency patterns that were suggestive of selection during domestication (**Supplementary Table 7**).

Fruit lycopene levels seem to have decreased upon the origin of SLC and then have largely remained in SLL (Razifard et al., 2020). We identified indels in genes that control lycopene biosynthesis and cyclization, of which the Heinz 1706 allele frequencies were significantly increased from SP to SLC and reached near fixation in heirlooms (**Supplementary Table 7**). These Heinz 1706 alleles, mostly deletions in promoters, were associated with decreased expression of *1-deoxy-D-xylulose 5-phosphate synthase* (*DXS*), *1-deoxy-D-xylulose-5-phosphate reductoisomerase* (*DXR*), *geranylgeranyl pyrophosphate synthase 2* (*GGPPS2*) and *ζ-carotene desaturase* (*ZDS*), and higher expression of *LCY-B* in the ripening fruit (**Fig. 3**), suggesting that these mutations might cause reduced lycopene levels in SLC and SLL by downregulating lycopene biosynthetic genes, and promoting lycopene degradation through upregulation of *LCY-B*. Interestingly, we found a recovery of certain LA2093 alleles in the elite tomato lines, including the ones potentially associated with higher expression of *DXR* and *GGPPS2* (**Supplementary Table 7**).

**Fig. 3.**
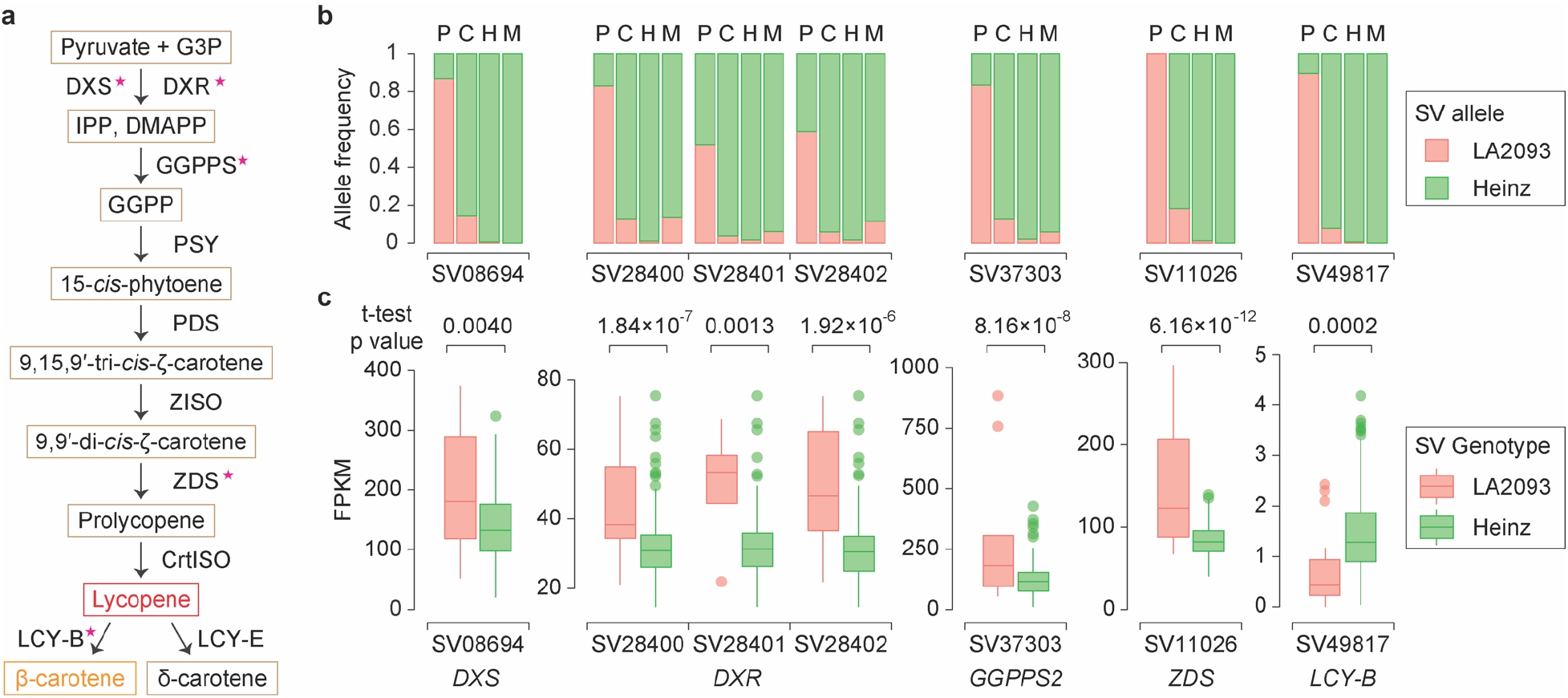
Selected SVs affecting the expression of lycopene metabolism genes. **a**, Lycopene biosynthesis and degradation pathway in tomato. Stars indicate enzyme-coding genes having selected SVs associated with significantly different gene expression levels between the two alleles (t-test p value <0.01). **b**, Allele frequencies of selected SVs in SP (P), SLC (C), heirloom (H) and modern (M) populations. **c**, Gene expression levels in tomato accessions carrying the homozygous LA2093 and Heinz 1706 alleles, respectively, of the selected SVs. For each box plot, the lower and upper bounds of the box indicate the first and third quartiles, respectively, and the center line indicates the median. G3P, glyceraldehyde 3-phosphate; IPP, isopentenyl diphosphate; DMAPP, dimethylallyl diphosphate; GGPP, geranylgeranyl diphosphate; DXS, 1-deoxy-d-xylulose 5-phosphate synthase; DXR, 1-deoxy-D-xylulose-5-phosphate reductoisomerase; GGPPS, geranylgeranyl pyrophosphate synthase; PSY, phytoene synthase; ZISO, ζ-carotene isomerase; ZDS, ζ-carotene desaturase; CrtISO, carotene isomerase; LCY-B, lycopene β-cyclase; LCY-E, lycopene ε-cyclase.

Fruit ripening regulation is of great interest to tomato researchers and breeders. Tomato FUL1, FUL2 and TAGL1 play key roles in controlling fruit ripening by interacting with RIN (Vrebalov et al., 2002; Bemer et al., 2012). Here, we identified a 15-bp in-frame deletion in *FUL1* of Heinz 1706, which had a frequency of 1.0%, 75.5% and 99.0% in SP, SLC and heirlooms, respectively. Intriguingly, *FUL1* was expressed at a considerably higher level in ripe fruits of the elite tomato line, NC EBR-1, carrying the derived deletion allele, than in LA2093 that carries the non-deletion allele (**Supplementary Fig. 10**). Consistently, accessions with homozygous *FUL1* deletion allele expressed the gene at a higher level in the orange-stage fruit than those with the non-deletion allele (**Supplementary Fig. 10**). Together, these results suggested that the *FUL1* deletion allele was associated with the higher *FUL1* expression and positively selected during domestication. *FUL2* of Heinz 1706 carried two deletions and one insertion in the promoter, all of which displayed allele frequency change patterns suggesting positive selection during tomato domestication and/or improvement and were associated with reduced expression of *FUL2* in the orange-stage fruit (**Supplementary Table 7**; **Supplementary Fig. 10**). The seemingly opposite effects of the selected alleles on the expression of *FUL1* and *FUL2* in orange-stage fruit could be related to the different spatial and temporal expression patterns of *FUL1* and *FUL2* in ripening fruit and their specific developmental functions (Bemer et al., 2012; Wang et al., 2014). In addition, two selected SVs in the introns of *TAGL1* were found to be associated with higher expression of *TAGL1* in cultivated tomatoes (**Supplementary Table 7**; **Supplementary Fig. 10**).

Wild species have been used as a source for improving SSC in cultivated tomatoes (Fridman et al., 2004; Petreikov et al., 2006). Three well-characterized genes that contribute to fruit SSC include *SUCR* controlling sucrose accumulation (Chetelat et al., 1995), and *Lin5* and *Agp-L1* regulating hexose content (Fridman et al., 2004). We found a 13-bp insertion in the first intron of *SUCR* associated with lower expression of the gene (**Supplementary Fig. 11**), whose frequency was 13.3% in SP, 86.5% in SLC, 99.4% in heirloom and fixed in modern accessions. LA2093 carried a 15-bp in-frame deletion in *Lin5*, resulting in a 5-aa deletion at one amino acid upstream of the critical amino acid for sucrose binding (Fridman et al., 2004). The frequency of the non-deletion allele was substantially increased during domestication (33.7% in SP, 90.9% in SLC, 98.9% in heirloom and fixed in modern accessions). For *AgpL1*, we identified an 18-bp deletion in the second intron and a 33-bp deletion in the promoter, both having significantly increased allele frequencies in SLC and SLL compared to SP (**Supplementary Table 7**). Domestication traits, including high fruit yield, increased fruit:leaf ratio and determinant habit, are found to be negatively correlated with high SSC (Chetelat et al., 1995). Although the Heinz 1706 alleles of these SSC genes were nearly fixed in the modern tomatoes, possibly a result of hitchhiking with domestication and improvement traits, the LA2093 alleles identified here offer an opportunity to improve SSC in the elite lines.

Incorporating alleles from wild species into SLL has been a strategy in tomato breeding to improve disease resistance. We identified selected SVs in a number of well-studied disease resistance genes (**Supplementary Table 7**). Further functional characterization of these SVs may open a door to recover the disease-resistance traits in cultivated tomatoes. We also identified selected SVs for genes involving in hormonal regulation, flower, inflorescence, seed and leaf development, as well metabolite biosynthesis (**Supplementary Table 7**), which would be associated with the dramatic changes of morphotype and metabolite diversity during the long history of tomato breeding.

### Genome-wide mapping of eQTLs

To explore the roles of SVs in gene expression regulation, we performed eQTL analysis using the published orange-stage fruit transcriptome data (Zhu et al., 2018). A total of 46,848 SVs in 10,789 eQTL regions were identified to be significantly associated with the expression of 5,595 genes, including 2,708 (25.1%) local and 8,081 (74.9%) distant eQTLs (**Fig. 4a** and **Supplementary Table 9)**. The local eQTLs were more significantly associated with the gene expression and thus explained more expression variation than the distant eQTLs (**Supplementary Fig. 12a, b**). The lead SVs of most local eQTLs were located near the start and end of coding regions (**Supplementary Fig. 12c**), indicating the important regulatory roles of sequences in these regions on gene expression.

**Fig. 4.**
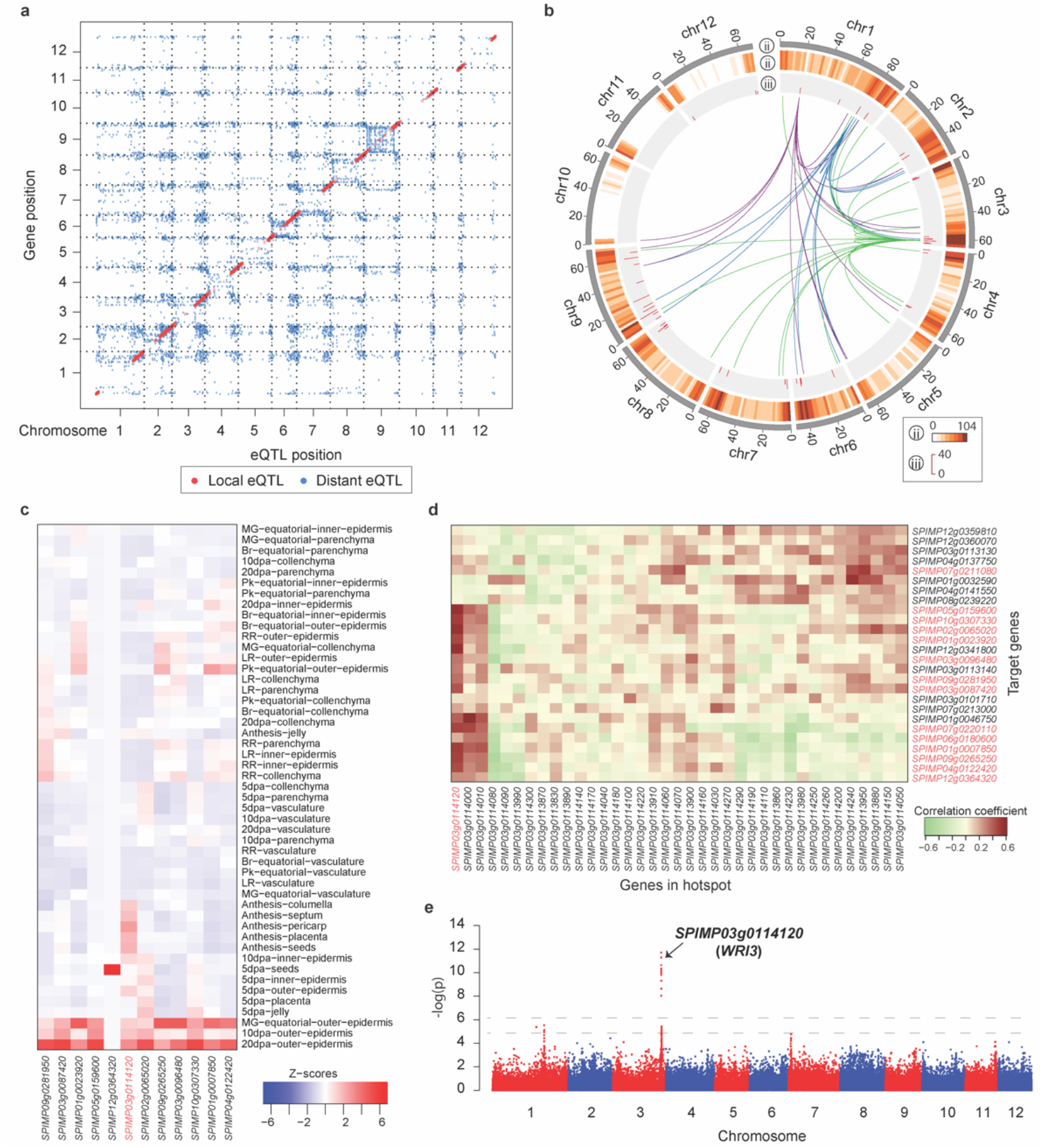
Genome-wide mapping of eQTLs. **a**, Positions of eQTLs identified in the genome. **b**, Distant eQTL hotspots. The outermost circle displays ideograms of the 12 tomato chromosomes. The second circle shows the number of target genes for all eQTLs in each 2-Mb window. The third circle shows the number of target genes of each distant hotspot. The innermost circle shows the links between three interesting eQTL hotspots and their target genes. Links between the three hotspots harboring the master regulators, A20/AN1 zinc finger protein, MYB12 and WRI3, respectively, and their target genes are depicted in purple, blue and green, respectively. **c**, Expression profiles of WRI3-targeted lipid biosynthetic genes in different tomato tissues. The *WRI3* gene (*SPIMP03g0114120*) is highlighted in red. **d**, Correlation coefficients between the expression levels of genes in the *WRI3*-hotspot and target genes. *WRI3* and the target lipid biosynthetic genes are highlighted in red. **e**, Manhattan plot of eQTLs associated with the *WRI3* expression. The horizontal dashed lines correspond to the Bonferroni-corrected significance thresholds at α = 0.05 and α = 1.

The genomic distribution of distant eQTLs was assessed to identify eQTL hotspots underlying the major regulatory variations across the tomato accessions. A total of 48 hotspots regulating the expression of 554 genes, as well as the potential master regulator within each hotspot, were identified (**Fig. 4b** and **Supplementary Table 10**). One hotspot located on chromosome 1 around 76.5 Mb (P=1.68×10^−17^) contained the *MYB12* gene, which encodes a key regulator of flavonoid biosynthesis (Adato et al., 2009; Ballester et al., 2010; Zhu et al., 2018). Among the 18 genes whose expression was regulated by this hotspot, seven encoded flavonoid biosynthetic enzymes (**Supplementary Table 11**), including chalcone synthase and flavonoid 3’-hydroxylase, which have been proved to be the direct targets of MYB12 (Zhang et al., 2015). Interestingly, another hotspot (Chr1:18800715-19894697; P =2×10^−11^) was found to affect the expression of the flavonoid biosynthetic genes regulated by the MYB12 hotspot. This hotspot contained an A20/AN1 zinc finger gene, whose homolog in poplar mediates flavonoid biosynthesis (Li et al., 2019). Furthermore, a total of 17 additional eQTLs were detected for 17 flavonoid biosynthetic genes (**Supplementary Fig. 13** and **Supplementary Table 12**), among which 14 genes had distant eQTLs and three had local eQTLs. Further study of these novel eQTLs will likely advance our knowledge on the gene regulatory network of flavonoid biosynthesis.

A major hotspot was found on chromosome 3 with 26 target genes, 14 of which were associated with lipid metabolism (**Supplementary Table 13**). An AP2/ERF transcription factor orthologous to Arabidopsis *WRI3* (To et al., 2012), was identified as the master regulator targeting 17 genes, including those encoding pyruvate kinase, pyruvate dehydrogenase, acetyl-coenzyme A carboxylase, carboxyl transferase and acyl carrier proteins (**Supplementary Table 13**). Co-expression of the tomato *WRI3* with lipid metabolism genes in the epidermis of developing fruit (Shinozaki et al., 2018) suggested its potential function in tissue-specific regulation of lipid biosynthesis, thus contributing to fruit cuticular wax accumulation (**Fig. 4c**). The expression of *WRI3* was positively correlated with that of all its target genes (**Fig. 4d**), suggesting its positive regulatory role. Furthermore, eQTL analysis identified two SVs, a 11-bp insertion in the sixth intron and a 77-bp deletion located in the protomer of *WRI3*, that were significantly associated with its expression (**Fig. 4e; Supplementary Fig. 14** and **Supplementary Table 7**). Altogether, these results suggested that *WRI3* could be a key regulator of fruit outer epidermis lipid deposition.

## Discussion

*S. pimpinellifolium* is thought to be the wild progenitor of cultivated tomatoes and offers desirable agronomic traits for breeding. Our study provides the first high-quality chromosome-scale SP reference genome that complements the existing tomato Heinz 1706 reference and serves as the foundation for exploring the genetic potential of SP and studying the genome evolution from wild to cultivated tomato under human selection. We performed explicit comparison between the genomes of LA2093 and Heinz 1706, resulting in a high-quality SV set further supported by evidence from the long-read split alignment. By investigating the frequency changes of wild alleles during tomato domestication, improvement and modern breeding, we not only revealed the population dynamics of known causal SVs underlying important traits, but also identified numerous novel candidate SVs in or near well-characterized genes controlling horticultural traits. The population dynamics of these novel SVs provide useful information for future determination of their likelihood of being the causal genetic variants. Furthermore, these SVs may be used as potential targets for future breeding programs to improve fruit quality and stress tolerance.

The LA2093 alleles of most selected SVs had significantly reduced frequencies after domestication and improvement. The massive loss of these wild alleles in cultivated tomatoes could be related to the large selective sweeps associated with domestication and improvement (Lin et al., 2014). Similar to the finding of metabolic genes hitchhiked with *fw11.3* (Zhu et al., 2018), linkage drag also contributed to the recovery of many LA2093 alleles in modern cultivars on chromosomes 4, 5 and 11, likely associated with recent intentional introduction of disease resistance, stress tolerance and favorable fruit quality traits into cultivars from wild accessions. Large structural variations complicate the cloning of genes underlying target traits and introgression of wild alleles in breeding without bringing along unwanted hitchhikers. Our study provides ample information that can be used to facilitate the recovery of wild beneficial alleles into modern elite cultivars.

eQTL analysis helped to unravel the hierarchical relationships among genes and provided insights into the effects of SVs on gene expression. In additional to confirming the central role of MYB12 in regulating flavonoid biosynthesis in tomato fruit, we identified additional novel eQTLs containing potential regulatory genes and SVs contributing to the complex regulatory network of flavonoid biosynthesis. Tomato fruit cuticular lipids provide a barrier against water loss and microbial infection (Isaacson et al., 2009; Martin and Rose, 2014). A novel AP2/ERF transcription factor gene, *WRI3*, may serve as a master regulator that controls the tissue-specific expression of key lipid biosynthetic enzyme genes in tomato fruit epidermis. These results increased our knowledge of the regulatory mechanism involved in fruit cuticular lipid accumulation, which could be applied to the development of crops with improved post-harvest performance.

## Methods

### Library construction and sequencing

Plants of the *Solanum pimpinellifolium* accession LA2093 were grown in the greenhouse at Boyce Thompson Institute in Ithaca, New York, with a 16-hour light period at 20°C (night) to 25°C (day). A PacBio SMRT library was constructed from the high molecular weight (HMW) DNA following the standard SMRTbell library preparation protocol and sequenced on a PacBio Sequel platform using the 2.0 chemistry (PacBio). Meantime, an Illumina paired-end library with insert size of ~450 bp was constructed using the Illumina Genomic DNA Sample Preparation kit following the manufacturer’s instructions (Illumina), and a Hi-C library was prepared following the proximo Hi-C plant protocol (Phase Genomics). Both the Illumina paired-end and the Hi-C libraries were sequenced on an Illumina NextSeq 500 platform with the paired-end mode and read length of 150 bp.

Nanopore sequences were generated for the SP accession LA1589 using the method described in Mazo-Molina et al. (2020) and used for validation of SV genotyping. Briefly, Nanopore libraries were constructed from the HMW DNA of LA1589 using the Ligation Sequencing Kit (SQK-LSK109) and sequenced on the MinION R9 flow cells for 48 h. Basecalling was performed using Guppy (v3.1.5).

Total RNA was extracted from immature, mature green and red ripe fruits using the QIAGEN RNeasy Plant Mini Kit (QIAGEN). Strand-specific RNA-Seq libraries were constructed using the protocol described in Zhong et al. (2011) and sequenced on an Illumina HiSeq 2000 platform (Illumina). Three biological replicates were performed for each sample.

### *De novo* assembly of the LA2093 genome

Raw PacBio reads were error corrected and assembled into contigs using CANU (Koren et al., 2017) (v1.7.1) with default parameters except that ‘OvlMerThreshold’ and ‘corOutCoverage’ were set to 500 and 200, respectively. PacBio reads were then aligned to the contigs and based on the alignments errors in the assembled contigs were corrected using the Arrow program implemented in SMRT-link-5.1 (PacBio). Furthermore, the Illumina paired-end reads were processed to remove adaptor and low-quality sequences using Trimmomatic (Bolger et al., 2014). The cleaned Illumina reads were aligned to the contigs using BWA-MEM (Li and Durbin, 2009) with default parameters, and based on the alignments two rounds of iterative error corrections were performed using Pilon (Walker et al., 2014) with parameters ‘--fix bases --diploid’. The final error-corrected contigs were then compared against the NCBI non-redundant nucleotide database, and those with more than 95% of their length similar to sequences of organelles (mitochondrion or chloroplast) or microorganisms (bacteria/fungi/viruses), were considered contaminants and discarded. The redundans pipeline (Pryszcz and Gabaldon, 2016) was then used to remove redundancies in the assembled contigs with parameters ‘--identity 0.99 --overlap 0.97’.

To scaffold the assembled contigs, Illumina reads from the Hi-C library were processed with Trimmomatic (Bolger et al., 2014) to remove adaptor and low-quality sequences. The cleaned Hi-C reads were aligned to the assembled contigs and the alignments were filtered using the Arima-HiC mapping pipeline (https://github.com/ArimaGenomics/mapping_pipeline). Based on the alignments, the contigs were clustered into pseudomolecules using SALSA (Ghurye et al., 2019) with parameters ‘-e GATC -i 3’. Furthermore, contigs of LA2093 were compared with the Heinz1706 reference genome (Hosmani et al., 2019) (version 4.0) using RaGOO (Alonge et al., 2019). Hi-C data and synteny with the Heinz1706 genome were integrated to construct the pseudomolecules. Finally, a genetic map constructed from a recombinant inbred line (RIL) population with LA2093 as one of the parents (Gonda et al., 2019) was used to validate the construction of pseudomolecules. Inconsistencies between the LA2093 pseudomolecules and genetic maps, and in the synteny with the Heinz1706 genome were manually checked and the accuracy of the LA2093 pseudomolecules was further validated using PacBio read alignment information.

### Annotation of the LA2093 genome assembly

MITE-Hunter (Han and Wessler, 2010) and LTRharvest (Ellinghaus et al., 2008) were used to *de novo* identify miniature inverted-repeat transposable elements (MITEs) and long terminal repeats (LTRs), respectively, in the assembled LA2093 genome. The LA2093 genome was masked using RepeatMasker (http://www.repeatmasker.org/) with the identified MITEs and LTRs, and the unmasked genome sequences were then analyzed using RepeatModeler (http://www.repeatmasker.org/RepeatModeler.html) to build a *de novo* repeat library. The final repeat library was obtained by combining the MITEs, LTRs and the *de novo* repeat library, and subsequently used to screen the LA2093 genome for repeat sequences using RepeatMasker. The identified repeat sequences were classified using the RepeatClassifier program of RepeatModeler.

Protein-coding genes were predicted from the repeat-masked LA2093 genome. LA2093 RNA-Seq reads generated in this study and from a previous study (Gonda et al., 2019) were processed to trim low-quality and adapter sequences using Trimmomatic (Bolger et al., 2014). The cleaned high-quality RNA-Seq reads were aligned to the assembled genome using HISAT2 (Kim et al., 2019) (v2.1) with default parameters. Transcripts were assembled from the read alignments using StringTie (Pertea et al., 2015) (v1.3.3b). The complete coding sequences (CDS) were predicted from the assembled transcripts using the PASA pipeline (Haas et al., 2003) (v2.3.3). *Ab initio* gene predictions were performed using BRAKER (Hoff et al., 2019), GeneMark-ET (Lomsadze et al., 2014), and SNAP (Korf, 2004). Protein sequences from the Swiss-Prot database and from Heinz 1706, *S. pennellii*, pepper and potato were used as protein homology evidence. Finally, the high-confidence gene models in the LA2093 genome were predicted using the Maker pipeline (Cantarel et al., 2008) by integrating *ab initio* predictions, transcript mapping and protein homology evidence.

### Detection of SVs and SNPs between reference genomes

To identity SVs between genomes of LA2093 and Heinz 1706 (SL4.0), the two genomes were first aligned using Minimap2 (Li, 2018) with the parameter ‘-ax asm5’. The resulting alignments were analyzed using Assemblytics (Nattestad and Schatz, 2016) for SV identification. The identified SVs spanning or close (<50 bp) to gap regions in either of the two genomes were excluded.

SVs were also identified using pbsv (https://github.com/PacificBiosciences/pbsv) and svim (Heller and Vingron, 2019) by aligning LA2093 PacBio reads to the Heinz1706 genome and Heinz 1706 PacBio reads to the LA2093 genome. The identified SVs spanning gap regions in the genomes were discarded. For large inversions identified through direct comparison of the two genomes, the two breakpoints of a candidate inversion were identified based on the genome alignment results. To confirm the breakpoints, the supported split reads were extracted from the results of pbsv and svim. Inversions were kept if more than 90% of total reads spanning the breakpoint were split reads or they were detected by both pbsv and svim. For the remaining SVs, the 5-kb flanking sequences of each SV were extracted from the reference genome, and then blasted against the query genome. The blast hits were then compared to the unique alignments between LA2093 and Heinz 1706 genomes identified by Assemblytics (Nattestad and Schatz, 2016). An SV was kept if the following criteria were met: 1) the blast hits of the two flank sequences of the SV (alignment length >50 bp, identity >90%, e-value <1e-10) were in the expected region on the query genome; and 2) the distance between the two blast hits was largely consistent with the SV size estimated by PacBio read mapping. For insertions, we required that the difference in SV size determined by PacBio read mapping and distance between the two blast hits was smaller than 20% of the estimated SV size. For deletions, the allowed gaps or overlaps between the two blast hits of flanking regions should be smaller than 3 bp. Repeat expansions/contractions and tandem expansions/contractions detected by Assemblytics (Nattestad and Schatz, 2016) were converted into one or more simple indels if the precise breakpoints were defined using the SVs identified by pbsv. SVs identified by both Assemblytics and pbsv were combined if they overlapped with each other (>50% in each). GO term enrichment analysis for the SV-related genes was performed using the Fisher’s exact test in the Blast2GO suite (Gotz et al., 2008) with a cutoff of adjusted P value <0.05.

SNPs between the two genomes were identified by comparing LA2093 and Heinz 1706 genomes using MUMmer4 (Marcais et al., 2018). The uniquely aligned fragments were used to identify SNPs with the show-snp tool in the MUMmer4 package.

### SV and SNP genotyping in the tomato population

Genome resequencing data of 725 tomato accessions reported in Gao et al (2019) were downloaded from the NCBI SRA database (**Supplementary Table 8**). The downloaded raw Illumina sequences from each accession were first processed to consolidated duplicated read pairs, which were defined as those having identical bases in the first 90 bp (for 100-bp reads) or 100 bp (for 150-bp reads) of both left and right reads, into unique read pairs. The resulting reads were processed to trim adaptor and low-quality sequences using Trimmomatic (Bolger et al., 2014).

The high-quality reference SVs identified between the LA2093 and Heinz1706 genomes were genotyped in the 725 tomato accessions. The cleaned Illumina reads from each accession were aligned to the LA2093 and Heinz1706 genomes, respectively, using BWA-MEM (Li and Durbin, 2009) allowing no more than 3% mismatches. For each SV in each accession, reads aligned to regions spanning the breakpoints of the SV in both LA2093 and Heinz1706 genomes were extracted and checked. For inversions, only split reads were used as evidence and each breakpoint was supported by at least three split reads. For indels with breakpoints supported by less than three split reads, the read coverage in the deleted regions were further checked. For a high-confidence indel, we required that <50% of the deleted region was covered by reads with 2× depth, while >50% of at least one of the flanking regions with the same length of the deleted region was covered. Based on the split read and read depth information, SVs in a particular accession were classified as the LA2093 genotype (same as in LA2093), the Heinz1706 genotype (same as in Heinz 1706), heterozygous (containing both LA2093 and Heinz1706 alleles), or undetermined (genotypes that could not be determined due to insufficient read mapping information). Accessions with less than 40% of SVs genotyped were excluded, leaving 597 tomato accessions kept for the downstream analyses.

Illumina reads aligned to the LA2093 genome were used for SNP calling. First, the duplicated alignments were marked using Picard (http://broadinstitute.github.io/picard/), with the parameter ‘OPTICAL_DUPLICATE_PIXEL_DISTANCE=250’. The ‘HaplotypeCaller’ function of GATK (McKenna et al., 2010) (version 3.8) was then used to generate a GVCF file for each accession with parameters ‘--genotyping_mode DISCOVERY --max_alternate_alleles 3 --read_filter OverclippedRead’, followed by the population variant calling using the function ‘GenotypeGVCFs’ with default parameters. Hard filtering was applied to the raw variant calling set, with parameters ‘QD<2.0 || FS> 60.0 || MQ< 40.0 || MQRankSum< −12.5 || ReadPosRankSum< −8.0’. Sites with at least 50% accessions genotyped and minor allele frequency (MAF) ≥0.03 and overlapping with the identified SNPs from the alignments of LA2093 and Heinz1706 genomes were kept.

### Population genomic analyses

Maximum-likelihood phylogenetic trees were constructed for the tomato accessions using the full SV dataset and SNPs at four-fold degenerated sites, respectively, using IQ-TREE (Nguyen et al., 2015) with 1000 bootstraps. Population structure was investigated using FastStructure (Raj et al., 2014) with default parameters. To analyze the selection of SVs during domestication, improvement and modern breeding, the frequency of each allele of a particular SV in each group was calculated. Significance of the difference of the frequencies between two compared groups was determined using Fisher’s exact test. The resulting raw P values of SVs were then corrected using the Bonferroni method and SVs with corrected P values <0.001 were defined as those under selection. The variation diversity for each chromosome and population differentiation index (*F*_ST_) across the genome were calculated using VCFtools (Danecek et al., 2011) (v0.1.16) based on the SVs and SNPs. *F*_ST_ values were calculated in each 1000-kb window with a step size of 250 kb.

Putative introgressions between two groups were identified using a likelihood ratio test (McNally et al., 2009) with the SV datasets. For each SV site in an accession of the heirloom or modern group, the percentage of accessions sharing this SV genotype was first derived in each of the two groups, then the ratio of the percentage of accessions sharing the genotype in the group this specific accession belonging to to that of accessions in the SP group was calculated. The average ratio for all SV sites in each of the 1000-kb windows with a step size of 250 kb were obtained. Regions with ratios of 0.9 or less and containing two or more SVs were defined as introgressions.

### eQTL analysis

Raw RNA-Seq data of 399 accessions reported in (Zhu et al., 2018) were downloaded from the NCBI SRA database under the accession SRP115430, of which 315 with SV data in this study were kept and used for the eQTL analysis. Raw sequencing reads were processed to trim low-quality and adaptor sequences using Trimmomatic (Bolger et al., 2014). The cleaned reads were aligned to the LA2093 genome using HISAT2 (Kim et al., 2019). Based on the alignments, raw read counts were derived for each gene and normalized to fragments per kilobase transcripts per million mapped fragments (FPKM). Genes with a median FPKM value of zero were excluded from the downstream analysis. Principal component analysis was performed based on the FPKM values, and nine accessions with FPKM values greater than 2.5 standard deviations from the mean in any of the first three principal components were excluded from the downstream analysis. To obtain a normal distribution of expression values for each gene, FPKM values of each gene were further normalized using the quantile-quantile normalization (qqnorm) function in R. To identify the hidden and confounding factors in the expression data, the normal quantile transformed expression values were processed using the Probabilistic Estimation of Expression Residuals (PEER) method (Stegle et al., 2012). The first 20 factors were selected as additional covariates in the genome-wide association studies. The Balding-Nichols kinship matrix constructed using EMMAX (Kang et al., 2010) with all SNPs and SVs was used to correct population structure. The missing genotypes in the raw biallelic SV dataset were imputed using the k-nearest neighbor (KNN) algorithm implemented in the fillGenotype software (Huang et al., 2010). In order to obtain the optimal imputation accuracy and filling rate, the accession with fewest missing genotypes in SP, SLC and Heirloom were selected and 10%, 20%, and 30% SV sites were randomly masked as missing genotypes for imputing. The imputations were performed using the fillGenotype with the combinations of the parameters: w = 20, 30, 50, 65 or 80), p = −3, −5, −7 or −9, k = 3, 5, 7 or 9), and r = 0.65, 0.7, 0.75 and 0.8. The optimal combination of parameters (w=80, k=3, p=−7, r=0.8) was selected after comparing the filling rate and imputation accuracy of each combination of the parameters (**Supplementary Table 14**). Only biallelic imputed SVs with minor allele frequency ≥ 1% and missing data rate ≤ 40% (a total of 71,684 SVs) were used for eQTL analysis. Genome-wide associations of transformed expression were estimated using the linear mixed model implemented in EMMAX (Kang et al., 2010). The Bonferroni test criteria at α = 0.05 and α = 1 were used as thresholds for significant and suggestive associations between variations and traits (expression), respectively, as described in (Li et al., 2012). In this study, the Bonferroni-corrected thresholds for the P values were 6.97×10^−7^ at α=0.05 and 1.40×10^−5^ at α=1, with corresponding −log10(P) values of 6.15 and 4.85, respectively.

Linkage disequilibrium (LD) decay was measured by calculating correlation coefficients (*r*^2^) for all pairs of SVs within 10 Mb using PopLDdecay (Zhang et al., 2019) (v3.27) with parameters ‘-MaxDist 500 -MAF 0.05 -Het 0.88 -Miss 0.999’. The stable *r*^2^ value was considered as the background level of LD. The background level of LD and physical distance were 0.205 and 1.94 Mb in this study, respectively. SVs within close vicinity to each other and associated with the expression of the same gene were grouped into an eQTL block, represented by the most significant SV (the lead SV) in the block. Two adjacent SVs were grouped into one region if the *r*^2^ >0.205 and physical distance <1.94 Mb, and regions with at least three significant SVs were considered as candidate eQTL blocks. eQTLs were considered local if they located within 50 kb of transcription start sites or transcription stop sites of the corresponding genes; otherwise, the eQTLs were considered distant.

eQTL hotspots were identified using the hot_scan software (Silva et al., 2014) with a window size of 50 kb and an adjusted P-value less than 0.05. Pairwise Pearson correlation coefficients between the target genes and genes located inside the eQTL hotspots were calculated using the FPKM values. The candidate master regulator was identified for each eQTL hotspot using the iterative group analysis (iGA) approach (Breitling et al., 2004).

## Data availability

The genome sequence of *Solanum pimpinellifolium* LA2093 has been deposited at DDBJ/ENA/GenBank under the accession JAAONP000000000. The version described in this paper is JAAONP010000000. The genome sequence of LA2093 and the associated annotations are also available at Sol Genomics Network (https://solgenomics.net/).

## ACKNOWLEDGMENTS

We thank Nabil Elrouby for collecting LA2093 RNA samples. This research was supported by grants from the US National Science Foundation (IOS-1855585 to Z.F. and J.J.G, IOS-1339287 to Z.F., J.J.G and C.C., and IOS-1546625 to G.B.M, Z.F. and S.R.S.).

## AUTHOR CONTRIBUTIONS

Z.F., S.W. and J.J.G. designed and managed the project. J.V, Z.F. and J.J.G. coordinated the LA2093 DNA extraction and genome sequencing. C.C. generated the LA2093 RNA-Seq data. J.Z., S.M. S.R.S and G.M performed the LA1589 Nanopore sequencing. P.H., S. Saha and L.A.M. contributed to the Heinz 1706 genome assembly and annotation. X.W., L.G., C.J., S.W. and S. Stravoravdis performed data analyses. X.W., L.G., S.W. and Z.F. wrote the manuscript. All authors reviewed and revised the manuscript.

## COMPETING INTERESTS

The authors declare no competing interests.

## Notes

### Competing Interest Statement

The authors have declared no competing interest.

## References

Adato, A., Mandel, T., Mintz-Oron, S., Venger, I., Levy, D., Yativ, M., Dominguez, E., Wang, Z., De Vos, R.C., Jetter, R., et al. (2009). Fruit-surface flavonoid accumulation in tomato is controlled by a SlMYB12-regulated transcriptional network. PLoS Genet 5, e1000777.

Alonge, M., Soyk, S., Ramakrishnan, S., Wang, X., Goodwin, S., Sedlazeck, F.J., Lippman, Z.B., and Schatz, M.C. (2019). RaGOO: fast and accurate reference-guided scaffolding of draft genomes. Genome Biol 20, 224.

Ashrafi, H., Kinkade, M., and Foolad, M.R. (2009). A new genetic linkage map of tomato based on a Solanum lycopersicum x S. pimpinellifolium RIL population displaying locations of candidate pathogen response genes. Genome 52, 935–956.

Ashrafi, H., Kinkade, M.P., Merk, H.L., and Foolad, M.R. (2012). Identification of novel quantitative trait loci for increased lycopene content and other fruit quality traits in a tomato recombinant inbred line population. Molecular Breeding 30, 549–567.

Ballester, A.R., Molthoff, J., de Vos, R., Hekkert, B., Orzaez, D., Fernandez-Moreno, J.P., Tripodi, P., Grandillo, S., Martin, C., Heldens, J., et al. (2010). Biochemical and molecular analysis of pink tomatoes: deregulated expression of the gene encoding transcription factor SlMYB12 leads to pink tomato fruit color. Plant Physiol 152, 71–84.

Bemer, M., Karlova, R., Ballester, A.R., Tikunov, Y.M., Bovy, A.G., Wolters-Arts, M., Rossetto Pde, B., Angenent, G.C., and de Maagd, R.A. (2012). The tomato FRUITFULL homologs TDR4/FUL1 and MBP7/FUL2 regulate ethylene-independent aspects of fruit ripening. Plant Cell 24, 4437–4451.

Blanca, J., Canizares, J., Cordero, L., Pascual, L., Diez, M.J., and Nuez, F. (2012). Variation revealed by SNP genotyping and morphology provides insight into the origin of the tomato. PLoS One 7, e48198.

Blanca, J., Montero-Pau, J., Sauvage, C., Bauchet, G., Illa, E., Diez, M.J., Francis, D., Causse, M., van der Knaap, E., and Canizares, J. (2015). Genomic variation in tomato, from wild ancestors to contemporary breeding accessions. BMC Genomics 16, 257.

Bolger, A.M., Lohse, M., and Usadel, B. (2014). Trimmomatic: a flexible trimmer for Illumina sequence data. Bioinformatics 30, 2114–2120.

Breitling, R., Amtmann, A., and Herzyk, P. (2004). Iterative Group Analysis (iGA): a simple tool to enhance sensitivity and facilitate interpretation of microarray experiments. BMC Bioinformatics 5, 34.

Cantarel, B.L., Korf, I., Robb, S.M., Parra, G., Ross, E., Moore, B., Holt, C., Sanchez Alvarado, A., and Yandell, M. (2008). MAKER: an easy-to-use annotation pipeline designed for emerging model organism genomes. Genome Res 18, 188–196.

Chen, G., Hackett, R., Walker, D., Taylor, A., Lin, Z., and Grierson, D. (2004). Identification of a specific isoform of tomato lipoxygenase (TomloxC) involved in the generation of fatty acid-derived flavor compounds. Plant Physiol 136, 2641–2651.

Chetelat, R.T., Deverna, J.W., and Bennett, A.B. (1995). Effects of the Lycopersicon chmielewskii sucrose accumulator gene (sucr) on fruit yield and quality parameters following introgression into tomato. Theor Appl Genet 91, 334–339.

Danecek, P., Auton, A., Abecasis, G., Albers, C.A., Banks, E., DePristo, M.A., Handsaker, R.E., Lunter, G., Marth, G.T., Sherry, S.T., et al. (2011). The variant call format and VCFtools. Bioinformatics 27, 2156–2158.

Ebert, A.W., and Schafleitner, R. (2015). Utilization of wild Relatives in the breeding of tomato and other major vegetables. In Crop Wild Relatives and Climate Change (John Wiley & Sons, Inc Hoboken, NJ, USA), pp. 141–172.

Ellinghaus, D., Kurtz, S., and Willhoeft, U. (2008). LTRharvest, an efficient and flexible software for de novo detection of LTR retrotransposons. BMC Bioinformatics 9, 18.

Foolad, M.R., Chen, F.Q., and Lin, G.Y. (1998). RFLP mapping of QTLs conferring cold tolerance during seed germination in an interspecific cross of tomato. Molecular Breeding 4, 519–529.

Fridman, E., Carrari, F., Liu, Y.S., Fernie, A.R., and Zamir, D. (2004). Zooming in on a quantitative trait for tomato yield using interspecific introgressions. Science 305, 1786–1789.

Gao, L., Gonda, I., Sun, H., Ma, Q., Bao, K., Tieman, D.M., Burzynski-Chang, E.A., Fish, T.L., Stromberg, K.A., Sacks, G.L., et al. (2019). The tomato pan-genome uncovers new genes and a rare allele regulating fruit flavor. Nat Genet 51, 1044–1051.

Gaut, B.S., Seymour, D.K., Liu, Q., and Zhou, Y. (2018). Demography and its effects on genomic variation in crop domestication. Nat Plants 4, 512–520.

Ghurye, J., Rhie, A., Walenz, B.P., Schmitt, A., Selvaraj, S., Pop, M., Phillippy, A.M., and Koren, S. (2019). Integrating Hi-C links with assembly graphs for chromosome-scale assembly. PLoS Comput Biol 15, e1007273.

Gonda, I., Ashrafi, H., Lyon, D.A., Strickler, S.R., Hulse-Kemp, A.M., Ma, Q., Sun, H., Stoffel, K., Powell, A.F., Futrell, S., et al. (2019). Sequencing-Based Bin Map Construction of a Tomato Mapping Population, Facilitating High-Resolution Quantitative Trait Loci Detection. Plant Genome 12.

Gotz, S., Garcia-Gomez, J.M., Terol, J., Williams, T.D., Nagaraj, S.H., Nueda, M.J., Robles, M., Talon, M., Dopazo, J., and Conesa, A. (2008). High-throughput functional annotation and data mining with the Blast2GO suite. Nucleic Acids Res 36, 3420–3435.

Guerrero, R.F., Rousset, F., and Kirkpatrick, M. (2012). Coalescent patterns for chromosomal inversions in divergent populations. Philos Trans R Soc Lond B Biol Sci 367, 430–438.

Haas, B.J., Delcher, A.L., Mount, S.M., Wortman, J.R., Smith, R.K., Jr., Hannick, L.I., Maiti, R., Ronning, C.M., Rusch, D.B., Town, C.D., et al. (2003). Improving the Arabidopsis genome annotation using maximal transcript alignment assemblies. Nucleic Acids Res 31, 5654–5666.

Han, Y., and Wessler, S.R. (2010). MITE-Hunter: a program for discovering miniature inverted-repeat transposable elements from genomic sequences. Nucleic Acids Res 38, e199.

Handsaker, R.E., Korn, J.M., Nemesh, J., and McCarroll, S.A. (2011). Discovery and genotyping of genome structural polymorphism by sequencing on a population scale. Nat Genet 43, 269–276.

Heller, D., and Vingron, M. (2019). SVIM: structural variant identification using mapped long reads. Bioinformatics 35, 2907–2915.

Hoff, K.J., Lomsadze, A., Borodovsky, M., and Stanke, M. (2019). Whole-Genome Annotation with BRAKER. Methods Mol Biol 1962, 65–95.

Hosmani, P.S., Flores-Gonzalez, M., van de Geest, H., Maumus, F., Bakker, L.V., Schijlen, E., van Haarst, J., Cordewener, J., Sanchez-Perez, G., Peters, S., et al. (2019). An improved de novo assembly and annotation of the tomato reference genome using single-molecule sequencing, Hi-C proximity ligation and optical maps. bioRxiv, 767764.

Huang, X., Wei, X., Sang, T., Zhao, Q., Feng, Q., Zhao, Y., Li, C., Zhu, C., Lu, T., Zhang, Z., et al. (2010). Genome-wide association studies of 14 agronomic traits in rice landraces. Nat Genet 42, 961–967.

Illa-Berenguer, E., Van Houten, J., Huang, Z., and van der Knaap, E. (2015). Rapid and reliable identification of tomato fruit weight and locule number loci by QTL-seq. Theoretical and Applied Genetics.

Isaacson, T., Kosma, D.K., Matas, A.J., Buda, G.J., He, Y., Yu, B., Pravitasari, A., Batteas, J.D., Stark, R.E., Jenks, M.A., et al. (2009). Cutin deficiency in the tomato fruit cuticle consistently affects resistance to microbial infection and biomechanical properties, but not transpirational water loss. Plant J 60, 363–377.

Kang, H.M., Sul, J.H., Service, S.K., Zaitlen, N.A., Kong, S.Y., Freimer, N.B., Sabatti, C., and Eskin, E. (2010). Variance component model to account for sample structure in genome-wide association studies. Nat Genet 42, 348–354.

Kim, D., Paggi, J.M., Park, C., Bennett, C., and Salzberg, S.L. (2019). Graph-based genome alignment and genotyping with HISAT2 and HISAT-genotype. Nat Biotechnol 37, 907–915.

Kinkade, M.P., and Foolad, M.R. (2013). Validation and fine mapping of lyc12.1, a QTL for increased tomato fruit lycopene content. Theor Appl Genet 126, 2163–2175.

Koren, S., Walenz, B.P., Berlin, K., Miller, J.R., Bergman, N.H., and Phillippy, A.M. (2017). Canu: scalable and accurate long-read assembly via adaptive k-mer weighting and repeat separation. Genome Res 27, 722–736.

Korf, I. (2004). Gene finding in novel genomes. BMC Bioinformatics 5, 59.

Li, H. (2018). Minimap2: pairwise alignment for nucleotide sequences. Bioinformatics 34, 3094–3100.

Li, H., and Durbin, R. (2009). Fast and accurate short read alignment with Burrows-Wheeler transform. Bioinformatics 25, 1754–1760.

Li, J., Sun, P., Xia, Y., Zheng, G., Sun, J., and Jia, H. (2019). A Stress-Associated Protein, PtSAP13, From Populus trichocarpa Provides Tolerance to Salt Stress. Int J Mol Sci 20.

Li, M.X., Yeung, J.M., Cherny, S.S., and Sham, P.C. (2012). Evaluating the effective numbers of independent tests and significant p-value thresholds in commercial genotyping arrays and public imputation reference datasets. Hum Genet 131, 747–756.

Lin, T., Zhu, G., Zhang, J., Xu, X., Yu, Q., Zheng, Z., Zhang, Z., Lun, Y., Li, S., Wang, X., et al. (2014). Genomic analyses provide insights into the history of tomato breeding. Nat Genet 46, 1220–1226.

Lomsadze, A., Burns, P.D., and Borodovsky, M. (2014). Integration of mapped RNA-Seq reads into automatic training of eukaryotic gene finding algorithm. Nucleic Acids Res 42, e119.

Marcais, G., Delcher, A.L., Phillippy, A.M., Coston, R., Salzberg, S.L., and Zimin, A. (2018). MUMmer4: A fast and versatile genome alignment system. PLoS Comput Biol 14, e1005944.

Martin, L.B., and Rose, J.K. (2014). There’s more than one way to skin a fruit: formation and functions of fruit cuticles. J Exp Bot 65, 4639–4651.

Mazo-Molina, C., Mainiero, S., Haefner, B.J., Bednarek, R., Zhang, J., Feder, A., Shi, K., Strickler, S.R., and Martin, G.B. (2020). Ptr1 evolved convergently with RPS2 and Mr5 to mediate recognition of AvrRpt2 in diverse solanaceous species. Plant J.

McKenna, A., Hanna, M., Banks, E., Sivachenko, A., Cibulskis, K., Kernytsky, A., Garimella, K., Altshuler, D., Gabriel, S., Daly, M., et al. (2010). The Genome Analysis Toolkit: a MapReduce framework for analyzing next-generation DNA sequencing data. Genome Res 20, 1297–1303.

McNally, K.L., Childs, K.L., Bohnert, R., Davidson, R.M., Zhao, K., Ulat, V.J., Zeller, G., Clark, R.M., Hoen, D.R., Bureau, T.E., et al. (2009). Genomewide SNP variation reveals relationships among landraces and modern varieties of rice. Proc Natl Acad Sci U S A 106, 12273–12278.

Mills, R.E., Walter, K., Stewart, C., Handsaker, R.E., Chen, K., Alkan, C., Abyzov, A., Yoon, S.C., Ye, K., Cheetham, R.K., et al. (2011). Mapping copy number variation by population-scale genome sequencing. Nature 470, 59–65.

Mu, Q., Huang, Z., Chakrabarti, M., Illa-Berenguer, E., Liu, X., Wang, Y., Ramos, A., and van der Knaap, E. (2017). Fruit weight is controlled by Cell Size Regulator encoding a novel protein that is expressed in maturing tomato fruits. PLoS Genet 13, e1006930.

Muller, N.A., Zhang, L., Koornneef, M., and Jimenez-Gomez, J.M. (2018). Mutations in EID1 and LNK2 caused light-conditional clock deceleration during tomato domestication. Proc Natl Acad Sci U S A 115, 7135–7140.

Nattestad, M., and Schatz, M.C. (2016). Assemblytics: a web analytics tool for the detection of variants from an assembly. Bioinformatics 32, 3021–3023.

Nguyen, L.T., Schmidt, H.A., von Haeseler, A., and Minh, B.Q. (2015). IQ-TREE: a fast and effective stochastic algorithm for estimating maximum-likelihood phylogenies. Mol Biol Evol 32, 268–274.

Pertea, M., Pertea, G.M., Antonescu, C.M., Chang, T.C., Mendell, J.T., and Salzberg, S.L. (2015). StringTie enables improved reconstruction of a transcriptome from RNA-seq reads. Nat Biotechnol 33, 290–295.

Petreikov, M., Shen, S., Yeselson, Y., Levin, I., Bar, M., and Schaffer, A.A. (2006). Temporally extended gene expression of the ADP-Glc pyrophosphorylase large subunit (AgpL1) leads to increased enzyme activity in developing tomato fruit. Planta 224, 1465–1479.

Pryszcz, L.P., and Gabaldon, T. (2016). Redundans: an assembly pipeline for highly heterozygous genomes. Nucleic Acids Res 44, e113.

Raj, A., Stephens, M., and Pritchard, J.K. (2014). fastSTRUCTURE: variational inference of population structure in large SNP data sets. Genetics 197, 573–589.

Razali, R., Bougouffa, S., Morton, M.J.L., Lightfoot, D.J., Alam, I., Essack, M., Arold, S.T., Kamau, A.A., Schmockel, S.M., Pailles, Y., et al. (2018). The Genome Sequence of the Wild Tomato Solanum pimpinellifolium Provides Insights Into Salinity Tolerance. Front Plant Sci 9, 1402.

Razifard, H., Ramos, A., Della Valle, A.L., Bodary, C., Goetz, E., Manser, E.J., Li, X., Zhang, L., Visa, S., Tieman, D., et al. (2020). Genomic Evidence for Complex Domestication History of the Cultivated Tomato in Latin America. Mol Biol Evol 37, 1118–1132.

Shen, J., Tieman, D., Jones, J.B., Taylor, M.G., Schmelz, E., Huffaker, A., Bies, D., Chen, K., and Klee, H.J. (2014). A 13-lipoxygenase, TomloxC, is essential for synthesis of C5 flavour volatiles in tomato. J Exp Bot 65, 419–428.

Shinozaki, Y., Nicolas, P., Fernandez-Pozo, N., Ma, Q., Evanich, D.J., Shi, Y., Xu, Y., Zheng, Y., Snyder, S.I., Martin, L.B.B., et al. (2018). High-resolution spatiotemporal transcriptome mapping of tomato fruit development and ripening. Nat Commun 9, 364.

Silva, I.T., Rosales, R.A., Holanda, A.J., Nussenzweig, M.C., and Jankovic, M. (2014). Identification of chromosomal translocation hotspots via scan statistics. Bioinformatics 30, 2551–2558.

Simao, F.A., Waterhouse, R.M., Ioannidis, P., Kriventseva, E.V., and Zdobnov, E.M. (2015). BUSCO: assessing genome assembly and annotation completeness with single-copy orthologs. Bioinformatics 31, 3210–3212.

Soyk, S., Lemmon, Z.H., Sedlazeck, F.J., Jimenez-Gomez, J.M., Alonge, M., Hutton, S.F., Van Eck, J., Schatz, M.C., and Lippman, Z.B. (2019). Duplication of a domestication locus neutralized a cryptic variant that caused a breeding barrier in tomato. Nat Plants 5, 471–479.

Stegle, O., Parts, L., Piipari, M., Winn, J., and Durbin, R. (2012). Using probabilistic estimation of expression residuals (PEER) to obtain increased power and interpretability of gene expression analyses. Nat Protoc 7, 500–507.

Su, X., Zhu, G., Huang, Z., Wang, X., Guo, Y., Li, B., Du, Y., Yang, W., and Gao, J. (2019). Fine mapping and molecular marker development of the Sm gene conferring resistance to gray leaf spot (Stemphylium spp.) in tomato. Theor Appl Genet 132, 871–882.

To, A., Joubes, J., Barthole, G., Lecureuil, A., Scagnelli, A., Jasinski, S., Lepiniec, L., and Baud, S. (2012). WRINKLED transcription factors orchestrate tissue-specific regulation of fatty acid biosynthesis in Arabidopsis. Plant Cell 24, 5007–5023.

Tomato Genome Consortium. (2012). The tomato genome sequence provides insights into fleshy fruit evolution. Nature 485, 635–641.

Vrebalov, J., Ruezinsky, D., Padmanabhan, V., White, R., Medrano, D., Drake, R., Schuch, W., and Giovannoni, J. (2002). A MADS-box gene necessary for fruit ripening at the tomato ripening-inhibitor (rin) locus. Science 296, 343–346.

Walker, B.J., Abeel, T., Shea, T., Priest, M., Abouelliel, A., Sakthikumar, S., Cuomo, C.A., Zeng, Q., Wortman, J., Young, S.K., et al. (2014). Pilon: an integrated tool for comprehensive microbial variant detection and genome assembly improvement. PLoS One 9, e112963.

Wang, S., Lu, G., Hou, Z., Luo, Z., Wang, T., Li, H., Zhang, J., and Ye, Z. (2014). Members of the tomato FRUITFULL MADS-box family regulate style abscission and fruit ripening. J Exp Bot 65, 3005–3014.

Xu, C., Liberatore, K.L., MacAlister, C.A., Huang, Z., Chu, Y.H., Jiang, K., Brooks, C., Ogawa-Ohnishi, M., Xiong, G., Pauly, M., et al. (2015). A cascade of arabinosyltransferases controls shoot meristem size in tomato. Nat Genet 47, 784–792.

Yuste-Lisbona, F.J., Fernandez-Lozano, A., Pineda, B., Bretones, S., Ortiz-Atienza, A., Garcia-Sogo, B., Muller, N.A., Angosto, T., Capel, J., Moreno, V., et al. (2020). ENO regulates tomato fruit size through the floral meristem development network. Proc Natl Acad Sci U S A.

Zhang, C., Dong, S.S., Xu, J.Y., He, W.M., and Yang, T.L. (2019). PopLDdecay: a fast and effective tool for linkage disequilibrium decay analysis based on variant call format files. Bioinformatics 35, 1786–1788.

Zhang, Y., Butelli, E., Alseekh, S., Tohge, T., Rallapalli, G., Luo, J., Kawar, P.G., Hill, L., Santino, A., Fernie, A.R., et al. (2015). Multi-level engineering facilitates the production of phenylpropanoid compounds in tomato. Nat Commun 6, 8635.

Zhong, S., Joung, J.G., Zheng, Y., Chen, Y.R., Liu, B., Shao, Y., Xiang, J.Z., Fei, Z., and Giovannoni, J.J. (2011). High-throughput illumina strand-specific RNA sequencing library preparation. Cold Spring Harb Protoc 2011, 940–949.

Zhu, G., Wang, S., Huang, Z., Zhang, S., Liao, Q., Zhang, C., Lin, T., Qin, M., Peng, M., Yang, C., et al. (2018). Rewiring of the Fruit Metabolome in Tomato Breeding. Cell 172, 249–261 e212.

